# Microbiota Assembly, Structure, and Dynamics Among Tsimane Horticulturalists of the Bolivian Amazon

**DOI:** 10.1101/779074

**Authors:** Daniel D. Sprockett, Melanie Martin, Elizabeth K. Costello, Adam Burns, Susan P. Holmes, Michael Gurven, David A. Relman

## Abstract

Little is known about the relative contributions of selective and neutral forces on human-associated microbiota assembly. Here, we characterize microbial community assembly in 52 Tsimane infant-mother pairs, using longitudinally collected stool and tongue swab samples profiled with 16S rRNA gene amplicon sequencing. The Tsimane are an indigenous Bolivian population who practice infant care associated behaviors expected to increase mother-infant dispersal. Infant consumption of dairy products, vegetables, and chicha (a fermented drink inoculated with oral microbes) was significantly associated with gut microbiota composition. At both body sites, maternal microbes at higher relative abundance were more likely to be shared. Shared microbes were also higher in abundance in infants at both body sites, but decreased in average relative abundance with age and were not significantly higher by 12 months of age. Infant microbiotas were modeled using a neutral community model of assembly, which showed that the prevalence of more than two thirds of infant-colonizing microbes could be explained using neutral processes alone. The same method was applied to datasets from Finnish and Bangladeshi infants, confirming that the majority of microbes colonizing infants from different countries were neutrally distributed. Among the Tsimane infant and adult gut microbiota samples, neutral processes were less prominent in villages with more market access. These results underscore the importance of neutral processes during infant microbiota assembly, and suggest that cultural changes associated with market integration may be affecting traditional modes of microbiota assembly by decreasing the role of these neutral processes, perhaps through changes in diet, sanitation, or access to medical care.

---

The human body becomes colonized in early life by microbial communities that are involved in several key metabolic and immunologic processes, including nutrient acquisition, immune programming, and pathogen exclusion. While these communities are relatively simple at birth, they grow in complexity until reaching an adult-like configuration within the first few years of life^1, 2^. This rapid and dynamic period of community assembly is characterized by broad shifts in community structure^3^ that arise from both deterministic processes like host-driven ecological selection^4^, as well as neutral processes like dispersal and demographic stochasticity^5–7^. Yet, the relative contributions of deterministic and neutral processes to early life microbiota assembly—and the extent to which different environmental factors may moderate these effects—remain largely unexplored.

In order to assess how transmission dynamics might affect community assembly, we focused on the Tsimane people, an indigenous forager-horticulturalist population inhabiting the Bolivian Amazon basin^8^. The Tsimane practice several infant care-associated behaviors (ICABs) that potentially increase the likelihood that maternally-derived microbes disperse to their offspring. For example, all Tsimane infants are vaginally birthed at home, and mothers carry infants in slings during the day and share their beds with their infants at night. Infants are breastfed ‘on demand’ 24 hours a day, and the period of exclusive breastfeeding lasts about 4 months^9, 10^, although typically, weaning is not complete until around 27 months of age^9^. During the introduction of complementary foods, mothers frequently premasticate, or pre-chew, foods such as rice, plantain, meat, or fish before depositing them into the mouths of their children. The most commonly consumed liquids are water (usually sourced from a nearby river), water mixed with sugar or fruit juice^11^, and chicha, a fermented drink made from manioc, corn, or plantain. Chicha made from manioc and corn is inoculated with saliva; women chew and expectorate pieces of manioc during preparation, and serve the drink without cooking. Similar chicha preparations have been shown to contain high loads of diverse lactic acid bacteria and yeast species^12–16^.

In comparison with industrialized societies, microbial exposure is high among the Tsimane. They lack access to improved water sources and have frequent exposure to parasites with high rates of early life morbidity and mortality owing to infectious disease^17–19^. Previous studies have reported that Tsimane children have high levels of C-reactive protein and other inflammatory markers, consistent with a high pathogen burden^19–22^. This confluence of high exposure to maternal and environmental sources of microbes and high immune system activation suggests high rates of microbial dispersal to Tsimane infants from family and environmental sources.

In recent years, rapid lifestyle changes have taken place in the Tsimane population due to increased access to market goods, cash economies, wage labor, education, and medicine^8^. These changes have not yet affected ICABs^10^, and the Tsimane continue to subsist primarily on horticulture and foraging, remain highly active, and exhibit negligible cardiometabolic health risks^23^. However, secular trends have documented increasing sedentarism, BMI, LDL cholesterol, and consumption of sugar, vegetable oil, and processed foods—particularly for Tsimane residing closer to market towns^23–25^. Thus, the Tsimane represent an important population in which to examine how changing lifestyle factors may affect patterns and processes of host-associated microbiota assembly.

Here, we report on microbiota data that we generated as part of a long-term study of Tsimane health and life history^8^. We first examined microbiota structure from paired fecal (n = 288) and oral (n = 119) samples collected longitudinally from 52 Tsimane mother-infant dyads between 2012-2013, one of the largest sample collections of this type to date from a non-industrialized setting. We then placed the patterns of microbiota assembly that we observed into a global context by comparing Tsimane infants to previously studied infants from Bangladesh^26^ and Finland^27^. Finally, we expanded our cohort by including data we generated from fecal samples collected in 2009 from 73 Tsimane individuals ranging in age from 1 to 58 years, enabling us to examine temporal and regional variation in microbiota composition associated with market integration. We found that even though neutral processes could largely explain patterns of infant microbiota assembly, these patterns were also consistent with an effect of ICABs. Gut bacterial taxa were associated with increasing market integration, and displayed altered dispersal patterns during infant colonization.

## RESULTS

To explore how ICABs affect patterns of microbial colonization in young children, stool samples (reflecting the distal gut) and swab samples from the dorsum of the tongue were collected from 52 Tsimane families living in six villages located along the Maniqui River in the Bolivian lowlands of the Amazon basin. Samples were collected from 48 infants (0-2 years of age) and 51 mothers (14 or more years of age) using a mixed longitudinal design (**Supplementary Table 1** and **Fig. 1a**). Bacterial communities in these samples were profiled using 16S rRNA gene amplicon sequencing. Anthropometric measures, as well as health and diet surveys, were also collected at the time of sampling^8^.

**Figure 1.**
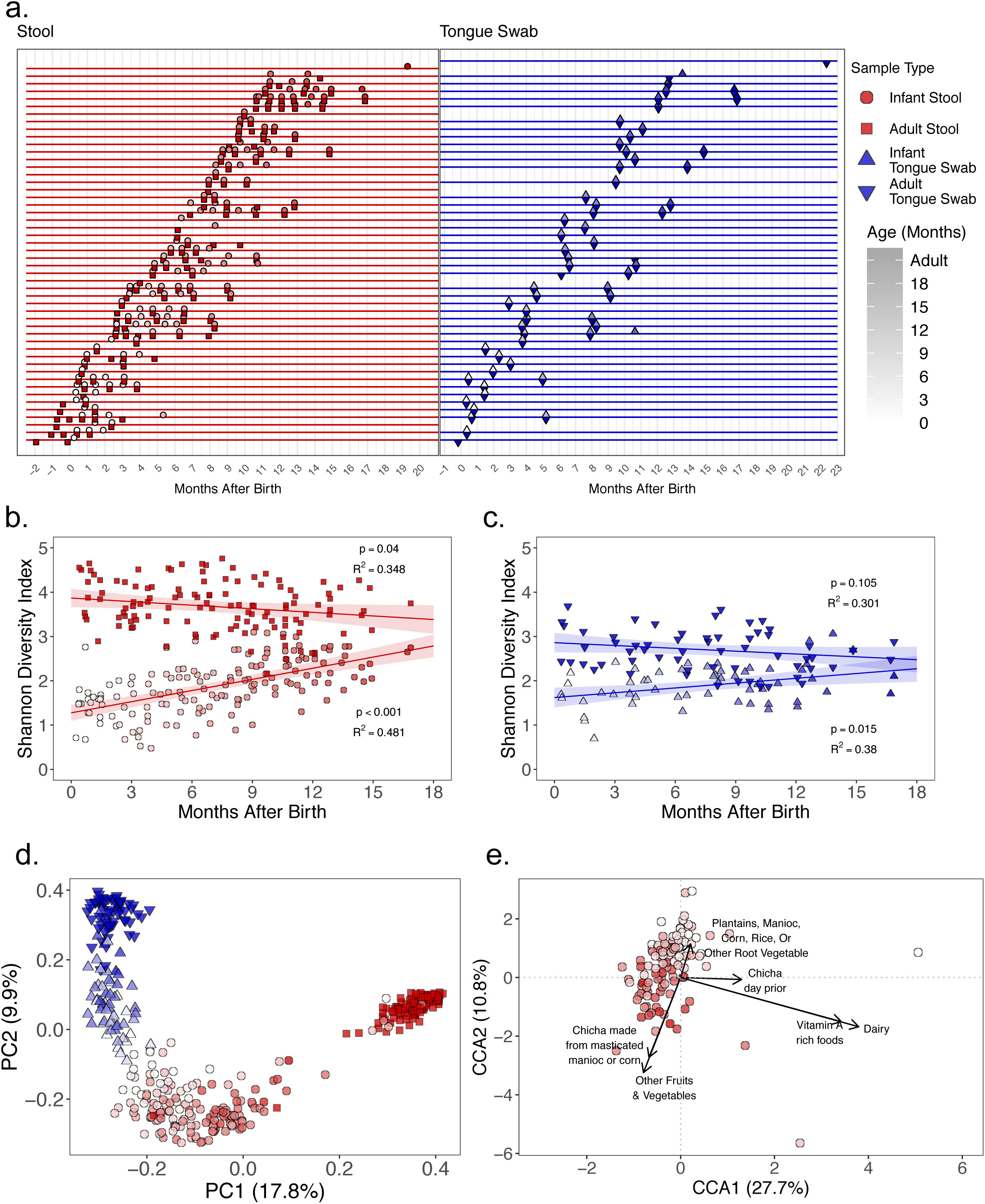
Infant microbiota dynamics and dietary factors associated with maturation. (a) Timeline denoting fecal sample and tongue swab collections for each mother-child dyad relative to the infant’s birth date. For maternal samples, time refers to the immediate postpartum period. Infant stool samples are circles, adult stool samples are squares, infant and adult tongue swabs are triangles that point up and down, respectively. Shapes are colored by sample type (red = stool, blue = tongue swabs), and the color darkens as the subject’s age increases. Shapes and colors are consistent across Figs. 1a–e. (b) Shannon Diversity Index regressed against the time since the infant’s birth for stool samples. Lines indicate the linear mixed-effects regression of diversity on time since delivery, while treating subject as a random effect. The shading indicates the 95% confidence interval. The conditional R^2^ describes the proportion of variation explained by both the fixed and random factors. (c) Shannon Diversity Index regressed against the time since the infant’s birth for tongue swab samples. Figure details are the same as in Fig. 1b. (d) Principal coordinate analysis (PCoA) of microbiota taxa composition in stool samples and tongue swabs from Tsimane dyads. (e) A partial canonical correspondence analysis (CCA) of ASV abundances, constrained against a matrix of diet survey data. The effects of infant age and village were controlled using a conditioning matrix. Significance was assessed using an ANOVA-like permutation test with 999 permutations, and only significant (p < 0.05) factors are displayed.

### Infant and Maternal Microbiota Dynamics Following Birth

Bacterial diversity increased with age in both the distal gut and tongue dorsum of infants over the first 18 months of life (**Fig. 1b,c**). In a linear mixed-effects model that included the subject as a random effect, diversity was significantly positively correlated with age in both stool (p < 0.001, conditional R^2^ = 0.481) and tongue swabs (p = 0.015, conditional R^2^ = 0.38), although diversity increased at a faster rate in the stool samples (**Fig. 1b,c**; ANOVA, p = 0.004). Maternal stool samples decreased in diversity over the study period (**Fig. 1b**, p = 0.04, conditional R^2^ = 0.348), which is consistent with other studies of the gut microbiota of women in the early postpartum period^28^. The diversity of the maternal dorsal tongue communities was stable during this time (**Fig. 1c**, p = 0.105).

Variation in bacterial community composition among samples from mothers and infants was primarily explained by body site (14%, PERMANOVA, p < 0.001) and host age (11%, PERMANOVA, p < 0.001). Village type (proximal or distal to the regional market) was also a significant factor (PERMANOVA, p < 0.01), although it explained less than 0.5% of the variation. Microbial communities colonizing the infant gut and tongue were distinct at all ages and became increasingly differentiated with time (**Fig. 1d** and **Supplementary Fig. 1**), but were never as distinct as those of adults, owing largely to incomplete maturation of these infant distal gut microbiotas by 18 months of age. The tongue swabs of 16-18 month olds harbored microbial communities similar in composition to adult tongue swabs (**Fig. 1d**), which contrasts with the findings of a previous cross-sectional study showing that the oral microbiota of Tsimane infants (9-24 months old) was distinct from that of their mothers^29^. The number of amplicon sequence variants (ASVs) shared between body sites within infants was highest immediately after birth (33.3%), and then declined with time (**Supplementary Fig. 1**). However, the ASV overlap between body sites in 16-18 month olds was significantly higher than in adults (**Supplementary Fig. 1**, Wilcoxon rank-sum test, p = 0.03).

Previous studies have reported a high prevalence of stunting and underweight in Tsimane children older than 2 years, but low prevalence of undernutrition^30^. Within the current study cohort of 48 Tsimane children, only one subject consistently exhibited height-for-age (HAZ) and weight-for-age (WAZ) z-scores more than 2 SD below the mean. Five other subjects had HAZ or WAZ scores that crossed from greater than to less than −2 SD after one year of age. All infants in the longitudinal cohort were breastfeeding at all study time points. However, among non-exclusively breastfed infants over 6 months of age, 63.8% of samples were collected at times when the infant had a minimum dietary diversity (MDD) score of <4 (**Supplementary Fig. 2a**), indicating potential micronutrient inadequacy in their complementary foods^31^. MDD is based on the 24-hour dietary recall of 7 key food groups, including grains, roots and tubers; legumes and nuts; dairy products (milk, yogurt, cheese); flesh foods (meat, fish, poultry and liver/organ meats); eggs; vitamin-A rich fruits and vegetables; and other fruits and vegetables. The infants’ MDD scores were positively correlated with infant gut diversity (**Supplementary Fig. 2a**, p = 0.027, conditional R^2^ = 0.324), and accounted for a larger effect than age (β_MDD_ = 1.2 vs. β_Age(months)_ = 0.6, p < 0.05 for both) on infant gut diversity. Chicha consumption, as well as vegetable and dairy consumption, was significantly associated with gut microbiota composition, even after controlling for age and village (**Fig. 1e**, partial CCA, ANOVA with 999 permutations, p < 0.05). Interestingly, consuming chicha made from manioc or corn had an opposite association with the infant gut microbiota than did consuming manioc or corn itself. In contrast to dietary factors, fecal neopterin, a marker of interferon-activated monocytes and macrophages, and of intestinal inflammation^32^, was associated with significantly lower infant gut diversity, even when controlling for age (**Supplementary Fig. 2b**, p = 0.002, conditional R^2^ = 0.333).

### Sources of Infant-Colonizing Microbes

To identify potential sources of infant-colonizing microbes, we first quantified the overlap between infant and adult microbiotas. 84.3% of the ASVs found in all infant stool samples at all time points were also found in at least one adult stool sample, while 79.9% of infant oral ASVs were also found in at least one adult oral sample. On average, about half of the microbes colonizing an infant’s gut or tongue were also found in their mother’s samples from the same body site, and this percentage increased as the infant aged (**Supplementary Fig. 3a**, blue line plus green line). However, the vast majority of these shared microbes were also found in other adults living in the same village as the infant (**Supplementary Fig. 3a**, green line), reflecting a high degree of microbial sharing among nearby adults, similar to that observed in other indigenous populations^33^. Mothers from the same village shared significantly more gut microbes, but not more oral microbes, than did mothers from different villages (**Supplementary Fig. 3b**, Wilcoxon rank-sum test, p < 0.001), suggesting that village ecology might play a role in structuring the gut microbiota. About half of the microbes found in an infant’s gut were found in at least one of their mother’s stool samples (**Supplementary Fig. 3c**, purple line plus green line), and about half were not found in any of their mother’s samples (orange line), with very few infant gut microbes ever observed on their mother’s tongue (yellow line). The same was true for infant tongue microbes, of which the majority of infant oral ASVs were observed on their mother’s tongue (**Supplementary Fig. 3c**, yellow line) and very few were observed in their mother’s gut (orange line).

Even though they accounted for only about half of the taxa in the infant distal gut and tongue dorsum, the bacteria shared within a mother-infant dyad were found at higher relative abundances in both the infants and their mothers than non-shared taxa. This difference in average relative abundance was statistically significant until the infants reached about 12 months of age, after which the relative abundances of shared and non-shared taxa did not significantly differ (**Fig. 2a,c**). At both body sites, the frequency at which a given microbe was shared within a dyad was largely a function of its relative abundance in the mother. Microbes with an average maternal abundance >0.1% were shared within a dyad at the same rate as its prevalence, while microbes at 0.01% or lower abundance in mothers were almost never shared within a dyad, even if the ASV was highly prevalent in infants (**Fig. 2b,d**). Overall, microbes that were highly abundant in Tsimane mothers were more likely to be shared with their infants.

**Figure 2.**
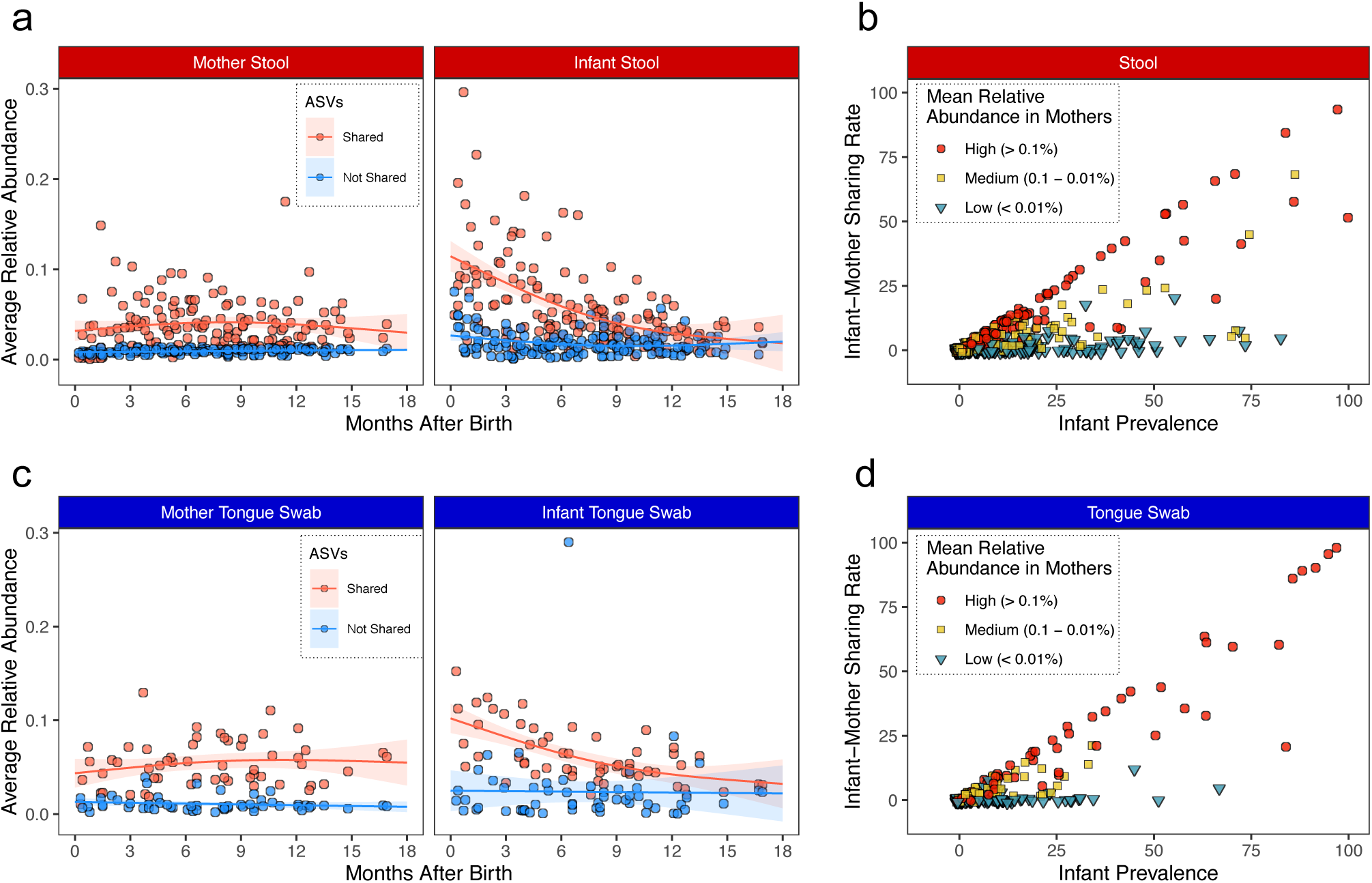
Shared microbes are highly abundant within mother-infant dyads. (a) Average relative abundance of ASVs that are shared or not shared within the stool samples of an infant-mother dyad plotted against the time since infant birth. Lines represent the best fit of a generalized additive model to either shared or not shared ASVs in the mother’s (left) or infant’s (right) stool samples. (b) Relationship between ASV prevalence in infant gut samples and the rate at which the ASV is shared within infant-mother dyads. The points are colored according to their average relative abundance in maternal gut samples. (c) Average relative abundance of ASVs that are shared or not shared within the tongue swabs of an infant-mother dyad plotted against the time since infant birth. Figure details are the same as in Fig. 2a. (d) Relationship between ASV prevalence in infant tongue swabs and the rate at which the ASV is shared within infant-mother dyads. Figure details are the same as in Fig. 2b.

### Microbiota Assembly Rules

Since many Tsimane ICABs may increase the opportunity for neutral dispersal of microbes from mothers to their children, we sought to quantify the relative contributions of deterministic and stochastic (*i.e.* neutral) processes during microbiota assembly. We used the neutral community model (NCM)^34^, which has previously been applied to the assembly of many host-associated microbial communities^6, 7^, to address this issue. The NCM predicts the prevalence of each microbe given its average relative abundance in a regional species pool. Microbes that fit those predictions are inferred to be assembling neutrally, while microbes at higher or lower prevalence are inferred to be under positive or negative ecological selection, respectively. The NCM assumes that all microbes in a regional species pool have an equal per capita ability to disperse to a local area, and once established, to have equal per capita fitness, growth, and death rates^34^.

The regional species pool was defined as containing all of the microbes able to disperse to and colonize a local area, which in this case was the infant gut or tongue. We estimated the composition of this pool by summing all of the local communities observed at a body site in all of the infants in the cohort (*i.e*., the ‘infant pool’). However, since we were also interested in the role that maternal microbes play in infant microbiota assembly, we assessed the degree to which the maternal microbiota contributed to the regional species pool by calculating the NCM goodness of fit (generalized R^2^) when maternal samples from the same body site were used instead to estimate the regional species pool (*i.e*., the ‘mother pool’). An R^2^ value close to 1 would imply that the composition of the infant gut microbiota was consistent with neutral dispersal, while a poor fit (*i.e*., R^2^ close to 0 or negative) would indicate that community assembly was due to more than just neutral dispersal. In this way, the NCM served as a useful mechanistic null model to help identify factors leading to a divergence from neutral dynamics or specific taxa that assembled in a non-neutral manner, such as strong competitors or active colonizers.

We found that microbes in the infant gut fit the neutral model moderately well when using the infant pool (**Fig. 3a**, R^2^ = 0.38), and that approximately two-thirds were neutrally distributed, while 27.9% experienced positive selection. Only 5.3% experienced negative selection; they included obligate anaerobes such as *Megasphaera*, *Prevotella*, *Bifidobacterium*, *Bacteroides,* and *Libanicoccus*, suggesting that oxygen exposure could be an impediment to transmission (**Fig. 3a**). Conversely, use of the mother pool resulted in a poorer model fit (R^2^ = − 0.29), indicating that infant-colonizing maternal microbes were under either positive or negative selection (**Fig. 3a**). Four microbes that were highly abundant and prevalent in the infant gut, *Escherichia/Shigella* sp. (ASV 6812), *Bifidobacterium* sp. (ASV 4271), *Veillonella atypica* or *V. dispar* (ASV 4083), and *Streptococcus* sp. (ASV 3537), were much less prevalent in the maternal gut (**Fig. 3b**). Conversely, two *Prevotella* spp. ASVs (5359 and 5371) were much less prevalent in the infant gut than the adult gut, especially in early life (**Fig. 3b**).

**Figure 3.**
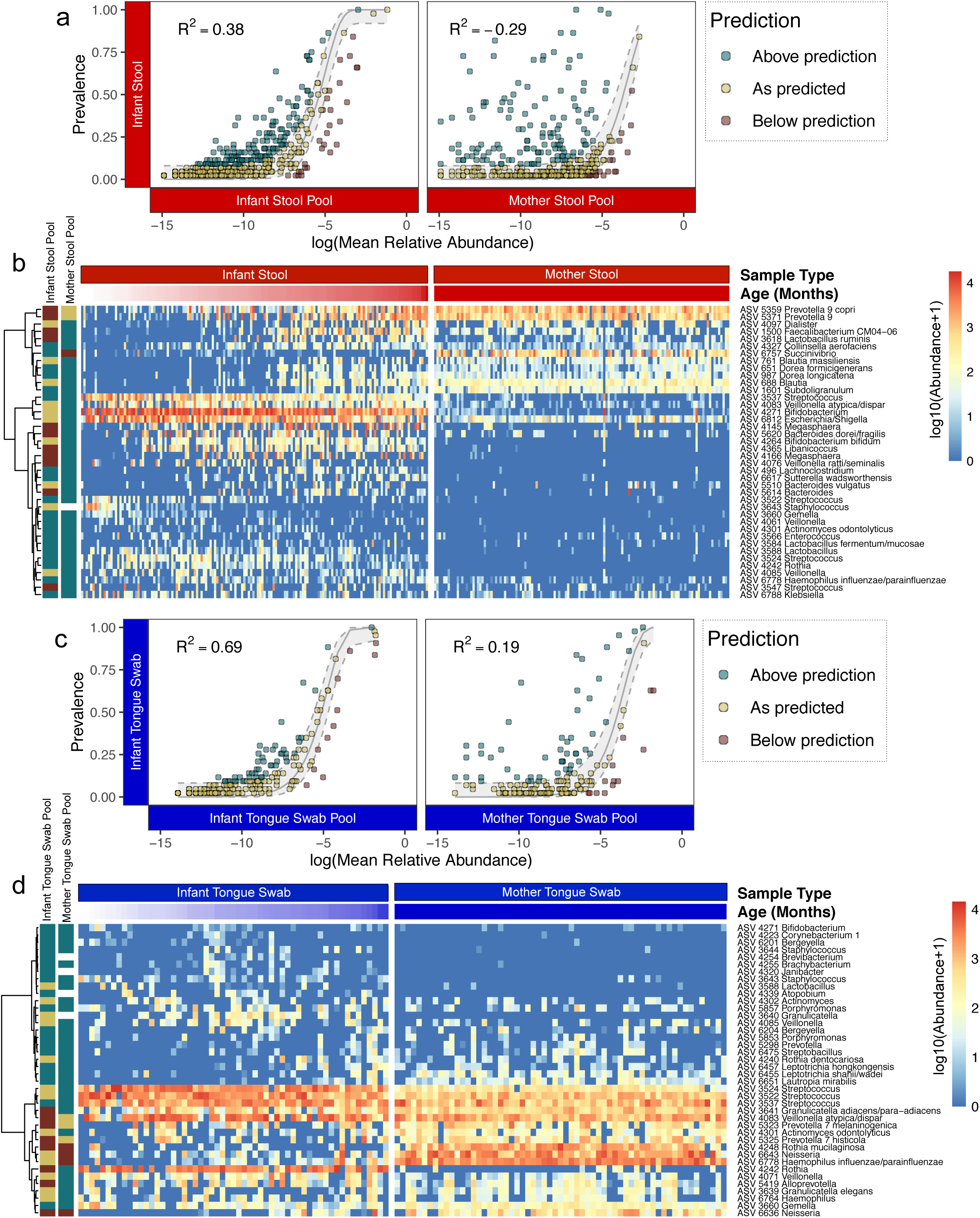
Infant microbiotas assemble according to neutral rules. (a) Neutral community model fit to each ASV observed in the stool of Tsimane infants, using either infant stool pool (left) or adult stool pool (right), as estimates of the regional species pool. Points are colored according to whether the taxon prevalence in infant samples is above (green), at (yellow), or below (red) the predicted prevalence according to the NCM. (b) Heatmap of the log normalized abundances of the ASVs from infant (left) and mother (right) stool samples that were observed in at least 25% of infant stool samples (n = 32). Rows were clustered on the y-axis using Ward’s minimum variance method. Row label colors correspond to the point colors in Fig. 3a. Columns were sorted by age for infants, and time since birth for mothers. (c) Neutral community model fit to each ASV observed from the tongue swabs of Tsimane infants, using either the infant tongue swab pool (left) or adult tongue swab pool (right), as estimates of the regional species pool. Figure details are the same as in Fig. 3a. (d) Heatmap of the log normalized abundances of ASVs from infant (left) and mother (right) stool samples that were observed in at least 25% of infant samples (n = 27). Figure details are the same as in Fig. 3b.

The NCM using the infant tongue swab pool predicted the distribution of infant oral microbes well, as the majority of these taxa assembled according to neutral expectations (Fig. 3c, R^2^ = 0.69). Interestingly, the model still fit relatively well using the mother tongue swab pool (Fig. 3c, R^2^ = 0.19), indicating that neutral mother-to-infant dispersal may play a more important role during assembly of the infant oral microbiota than the gut microbiota. *Streptococcus* spp. (ASVs 3524, 3522, and 3537), *Granulicatella adiacens* or *G. para-adiacens* (ASV 3641), and *Veillonella* sp. (ASV 4083) were all highly prevalent in both infant and mother tongue swabs. Two of these ASVs (4083 and 3537) were highly prevalent in the infant gut but not the adult gut, suggesting that a small number of bacterial taxa may disperse from the adult mouth to the infant gut and mouth.

### Infant Microbiota Assembly in a Global Context

We next sought to determine whether the assembly patterns that we observed in the Tsimane were conserved in infants from diverse cultures. We integrated our data with two publicly available datasets generated with comparable molecular methods for profiling the distal gut microbiota. The first of these datasets was created in a study of undernourished infants in Bangladesh^26^ (we used the well-nourished control group only), and the second in a large observational study of type 2 diabetes in eastern Europe^27^ (we used the Finnish infants only). Since the European study did not include samples from adults, we also included gut microbiota data from British adults to approximate the microbiota structure of a typical European, high-income country^35^. Despite drastically different social and physical environments, access to health care, and diets, the distal gut microbiota of the infants in these three studies were similar in composition at the earliest ages based on their Jaccard distance, indicating similar initial microbial colonists in societies across the globe (**Fig. 4a**). A principal curve was fit to each dataset, representing the average trajectory of microbiota composition across time in infants from that country. The age ranges for each infant cohort were 0.2 – 19.3 months for Bolivia (Tsimane), 0 – 24.2 months for Bangladesh, and 1.2 – 36.9 months for Finland. As the children became older, their microbiota trajectories diverged, and their microbiotas became more distinct as they approached a population-specific adult-like state (**Fig. 4a**). By scaling the principal curve position to the mean of the adult samples, and then regressing against the age of the subject we found that infant microbiotas from these three countries matured, or reached an average adult-like state, at significantly different rates (ANOVA, p > 0.05 for all pairwise comparisons) (**Supplementary Fig. 4**).

**Figure 4.**
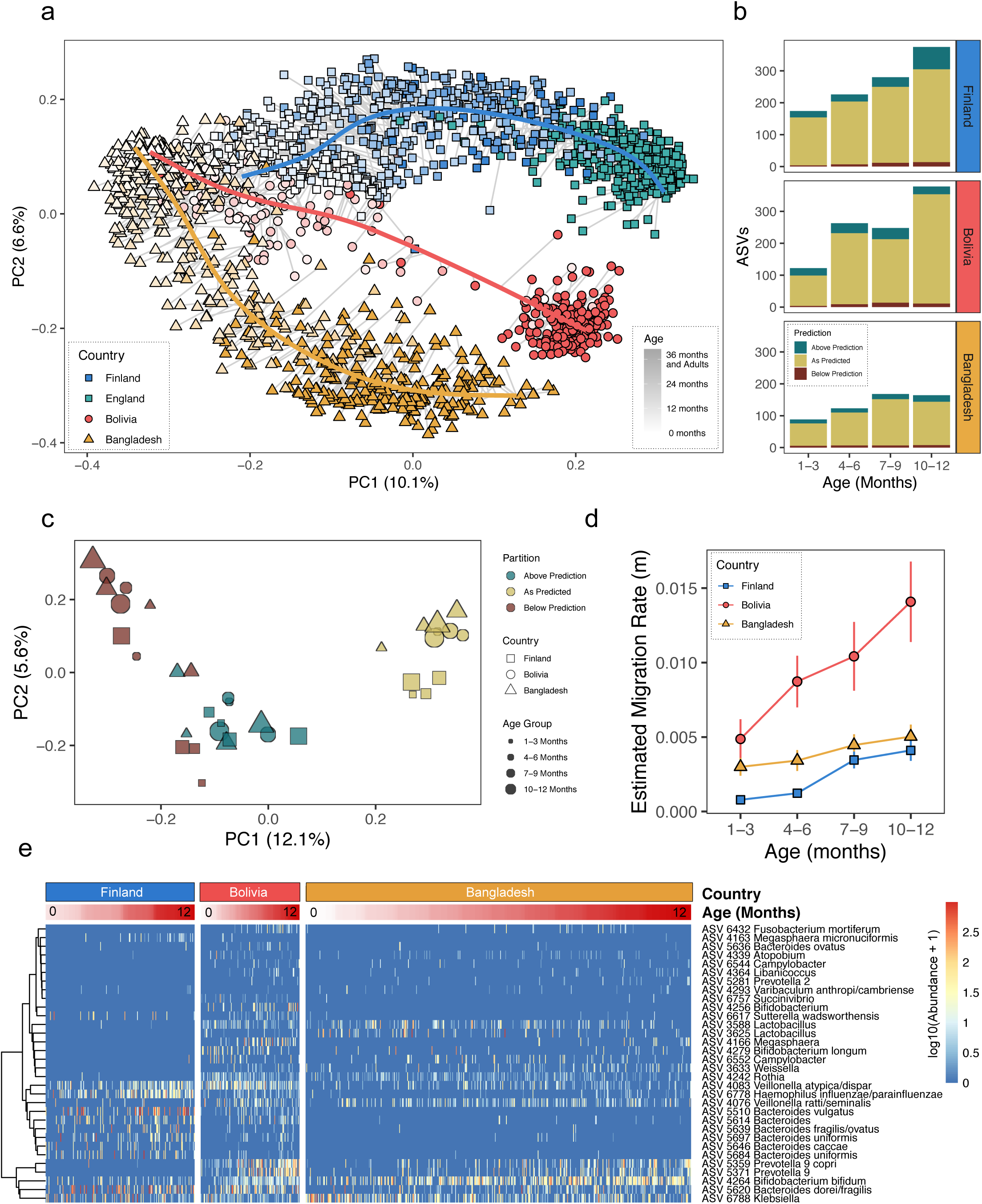
Patterns of selection and neutral assembly are consistent across ontogeny and geography. (a) Principal coordinate analysis (PCoA) of stool samples from Bolivian (Tsimane) dyads, as well as previously published 16S rRNA datasets of Bangladeshi and Finnish infants, as well as English adults. Points are shaped and colored according to country, and the color darkens with increased age of the subject (Finland - blue squares, England – green squares, Bolivia (Tsimane) – red circles, Bangladesh – yellow triangles). A principal curve was fit for each of the three populations (Bolivian (Tsimane), Bangladeshi, and European (Finnish/English)). (b) A summary of the proportion of ASVs present in each country and age group that fit the model (yellow), or was above (green) or below (red) the model’s prediction. See Supplementary Fig. 5 for details. (c) Principal coordinate analysis using the Jaccard distance for each of the 3 data partitions of four age groups from 3 countries (36 measurements in total). The color indicates the predicted fit, the shape indicates the country whence the samples were collected, and the size of each shape is scaled to indicate the age group of the subject. (d) The estimated migration rate (*m*) given by the neutral community model for each country and age group. Vertical lines represent 95% confidence intervals. Colors and shapes are the same as in Fig. 4a. (e) A heatmap of the log normalized abundances of ASVs that were estimated to be above their predicted prevalence given their average relative abundance in all 12 green partitions in Fig. 4c. Rows were clustered on the y-axis using Ward’s minimum variance method. Columns were sorted by infant age (0-12 months).

In order to determine if neutral processes operate in similar manners among infants in different cultural settings and geographies, we divided each dataset into three-month age intervals up to one year of age, and then refit the NCM model using only the samples from that interval as the estimate of the regional species pool, building a total of 12 additional models (**Supplementary Fig. 5**). These datasets exhibited striking consistency in the role of neutral and non-neutral processes across both age and geography, with the majority of microbes exhibiting patterns of neutral assembly across individuals (**Fig. 4b**). Furthermore, by partitioning ASVs according to country, age group, and how well they fit the neutral model, we found similar sets of neutral and non-neutral microbes across geography and ontogeny (**Fig. 4c**, Jaccard distance/PCoA). A heatmap of only the microbes that were positively selected revealed three, *Bifidobacterium bifidum* (ASV 4264), *Bacteroides dorei or B. fragilis* (ASV 5620), *and Klebsiella* sp. (ASV 6788) as common in all three cohorts (**Fig. 4e**). Other country-specific patterns were also apparent, including a set of 6 *Bacteroides* ASVs (ASVs 5510, 5614, 5639, 5697, 5646, and 5684) that were prevalent in Finnish and Bolivian (Tsimane), but not Bangladeshi infants, as well as two *Prevotella* ASVs (ASVs 5359 and 5371) that increased in prevalence with age in both the Bolivian (Tsimane) and Bangladeshi infants, while remaining nearly absent in Finnish infants (**Fig. 4e**). Interestingly, *Bacteroides* and *Prevotella* have been identified as biomarkers for Western and non-Western lifestyles^36^, respectively, yet both were prevalent in Tsimane infants.

The neutral community model also predicts the probability that when an individual microbe is removed from the local community, it is replaced by a microbe dispersing from the regional species pool rather than by reproduction from within the local community^7, 34^. This migration term, *m*, increased with age in all three populations, although it increased significantly more in the Tsimane infants (**Fig. 4d**). This significantly higher rate of microbial migration into the Tsimane infant gut is consistent with their high exposure to maternal and environmental microbes.

### Market Access Influences Tsimane Microbiota Structure

In recent decades, the Tsimane have experienced profound changes to their culture and life history, including increased access to processed foods in markets and greater access to modern medical care. In order to understand how these lifestyle shifts may have affected the assembly of the gut microbiota, we compared the gut microbiota structure of adults living in nine Tsimane villages. These villages belonged to three village ecotypes, ‘proximal river’, ‘distal river’, and ‘forest’, reflecting a gradient of cultural and lifestyle features currently in transition (**Supplementary Table 1** and **Fig. 5a**), such as higher intake of processed food^24^, increased obesity and metabolic syndrome^37^, and higher levels of urinary phthalates^38^, suggesting that increased economic integration and market access can have important health impacts. At the time of sample collection, proximal river villages (≤ 20 km to market) were accessible by road/taxi or boat, whereas distal river villages (>20 km to market) were only accessible by foot or boat (**Supplementary Table 1**). Samples from the two proximal and four distal river villages are those depicted in **Fig. 1a**. In addition, we included a single stool sample from each of 73 Tsimane individuals living in one of three forest villages (**Supplementary Table 1**). The set of samples from the river villages was collected in September 2012 through March 2013, while that from forest villages was collected in July 2009.

**Figure 5.**
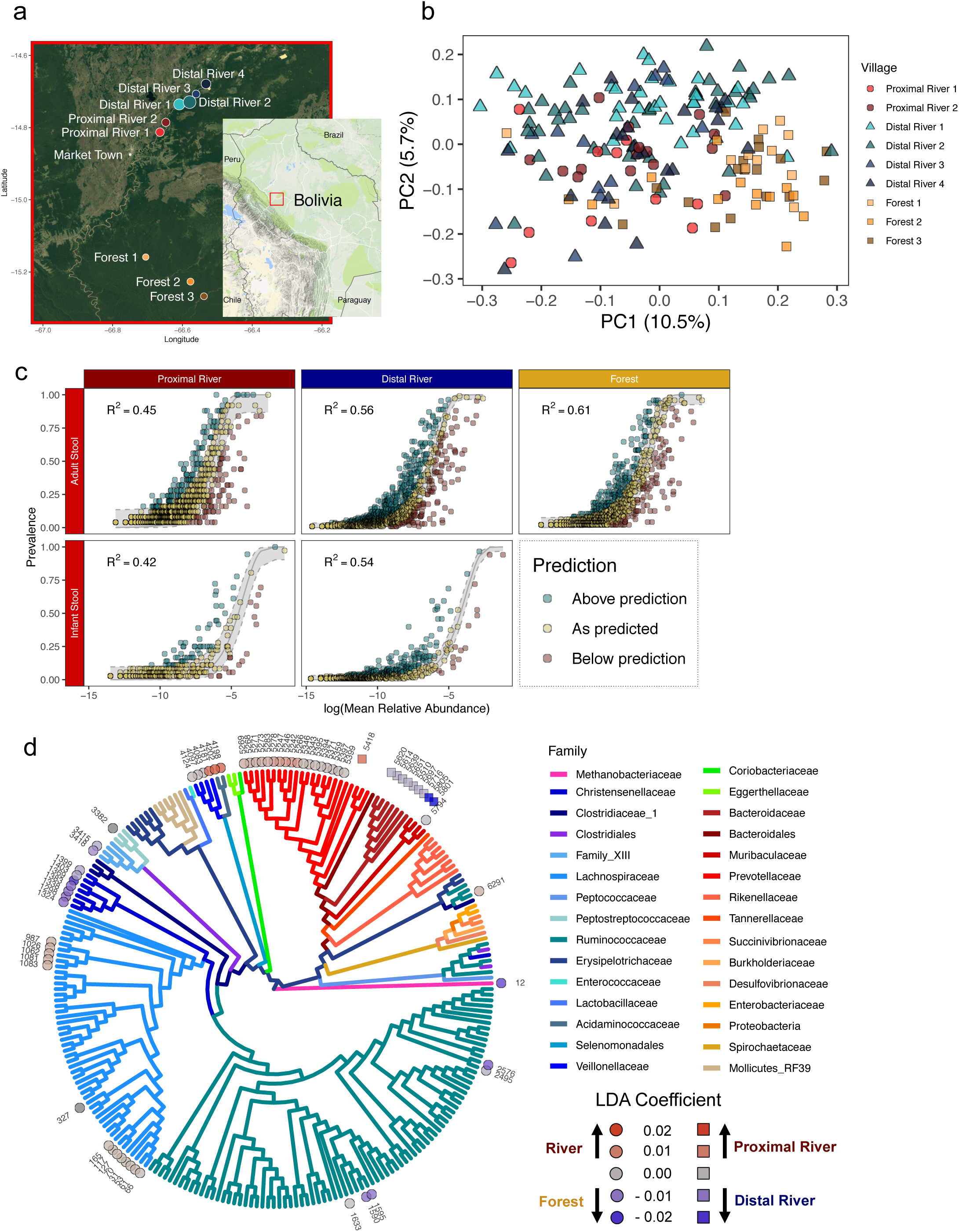
Market access is associated with Tsimane gut microbiota structure. (a) A map showing the location of the 9 Tsimane villages whence stool samples were collected. The size of each circle was scaled to the number of samples collected from that village, with the exception of the market town, where no samples were collected. The map insert denotes the region of Bolivia being displayed. (b) Principal coordinate analysis (PCoA) on the Bray-Curtis distance of 16S rRNA profiles from adult stool samples. Village types (‘proximal’ river, ‘distal’ river, and ‘forest’) have significantly different distributions (PERMANOVA with 1000 permutations, p = 0.001). Samples are colored by village and their shapes denote the village type (proximal river villages – circles, distal river villages – triangles, forest villages – squares). (c) Neutral community model fit to each ASV observed in the stool of Tsimane adults (top row) or infants (bottom row) from each of three village types (proximal river – left, distal river – right, and forest – left). In each case, the composition of the regional species pool was estimated using the samples from the age group and village type. The gray line is the predicted distribution (shaded area is 95% CI), and points are colored according to whether their prevalence in infant samples was above (green), at (yellow), or below (red) their predicted prevalence according to the NCM. (d) Phylogenetic tree with ASVs found at least 10 times in at least 20% of Tsimane adult stool samples. Discriminatory ASVs were identified using the tree-based LDA algorithm in the treeDA R package (version 0.0.3). ASVs are denoted by shapes based on two sets of comparisons (circles for river vs. forest and squares for proximal river vs. distal river), and are colored based on their discriminatory strength. ASVs are colored by the taxonomic family and significantly discriminatory ASVs are labeled with their ASV number.

A principal coordinates analysis of these microbiota data showed clustering of stool samples by ecotype of the village in which the subject lived (**Fig. 5b**, PERMANOVA, p < 0.05). PC1, which explained 10.5% of the variation in the data, was significantly correlated with the diversity of the microbiotas (**Supplementary Fig. 6**, p < 0.001, R^2^ = 0.289), although there was no significant difference in diversity between village types (p < 0.05 for all comparisons). To evaluate whether these relationships might reflect an effect of market access on assembly processes, we applied the NCM to samples from infants and adults living in the three different village ecotypes with the regional species pool estimated from all of the samples from that age group and village ecotype, and found that the goodness of fit decreased in samples from villages with higher levels of market access (**Fig. 5c**). This suggested that market integration, perhaps through increased access to processed foods (*i.e.* breads, pasta, refined sugar) and/or medicine (*i.e.* antibiotics), may increase the role of ecological selection and diminish the role of stochastic dispersal during assembly of the gut microbiota.

To identify how even subtle ecological and economic differences among villagers might affect the gut microbiota, we used a phylogeny-based implementation of linear discriminant analysis. This approach uses both the leaves of the phylogenetic tree (i.e., the ASVs), as well as the nodes to identify linear combinations of features that characterize two or more sample classes. The first model, built to discriminate between proximal and distal river villages, selected 3 predictors corresponding to 10 ASVs. Nine of those ASVs, including 2 in the family Muribaculaceae (ASVs 5801 and 5805) and 7 in the family Bacteroidaceae (ASVs 5614, 5639, 5697, 5510, 5620, 5651, 5716) were discriminatory for the distal river samples, while only 1 ASV, in the family Prevotellaceae (ASV 5418), was discriminatory for the proximal river samples (**Fig. 5d**). Even though diversity was not significantly lower in proximal river villages (**Supplementary Fig. 6**), the small number of proximal village-discriminatory features is consistent with studies suggesting that life in a high resource setting often leads to loss of gut bacterial diversity^39, 40^. Despite the small number of optimum features following 10-fold cross validation, the model accurately predicted village type for 87.3% of samples (**Supplementary Fig. 7**).

The second model, built to discriminate between river and forest villages, identified 24 predictors corresponding to 56 ASVs, and correctly identified the village type for 91.8% of the samples (**Supplementary Fig. 7**). River villages were associated with ASVs in the bacterial families Prevotellaceae (18 ASVs), Lachnospiraceae (13 ASVs), Veillonellaceae (4 ASVs), Acidaminococcaceae (2 ASVs), and Ruminococcaceae (2 ASVs). Indicator taxa for Tsimane forest villages included ASVs in the bacterial families Ruminococcaceae (4 ASVs), Clostridiales Family XIII (2 ASVs), Lachnospiraceae (1 ASV), Muribaculaceae (1 ASV), Peptostreptococcaceae (1 ASV), and Christensenellaceae (7 ASVs), as well as 1 archaeal ASV in the family Methanobacteriaceae (1 ASV) (**Fig. 5d**).

Christensenellaceae and Methanobacteriaceae were previously identified in a study of adult twins as members of a co-occurrence network of heritable gut microbiotas that were also associated with body leanness^35, 41^. Since Tsimane have low but increasing rates of obesity^23^, we sought to determine if a similar network was present in the Tsimane adults. The optimal network topology constructed from adult stool microbial communities contained a subnetwork involving the most highly connected node, ASV 12 (*Methanobrevibacter*), as well as 5 ASVs in family Christensenellaceae (**Fig. 6a**), indicating strong associations among these taxa. This subnetwork also included the second most highly connected node, *Prevotella copri* (ASV 5359). Although these two taxa frequently co-occurred, their abundances were inversely related (**Fig. 6b**, p < 0.001, marginal R^2^ = 0.205). *Methanobrevibacter and Prevotella* have been shown to be positively correlated in the guts of humans^42^ and western lowland gorillas^43^, especially in the setting of seasonal, carbohydrate-rich diets. These taxa also had distinct colonization dynamics in the gut of infants from Bolivia (Tsimane), Bangladesh, and Finland. *Prevotella copri*, the same ASV that was abundant in Bolivian and Bangladeshi infants but essentially absent in Finnish infants, increased in abundance beginning at approximately 6 months of age, and remained at high levels throughout adulthood (**Fig. 6c**). *Methanobrevibacter*, on the other hand, was not detected in these gut microbiotas until later in adolescence (**Fig. 6c**). However, since *Methanobrevibacter* was more common in forest villages, and these villages did not include many samples from young children, this pattern may also have been a result of ecological, seasonal, or societal differences between villages.

**Figure 6.**
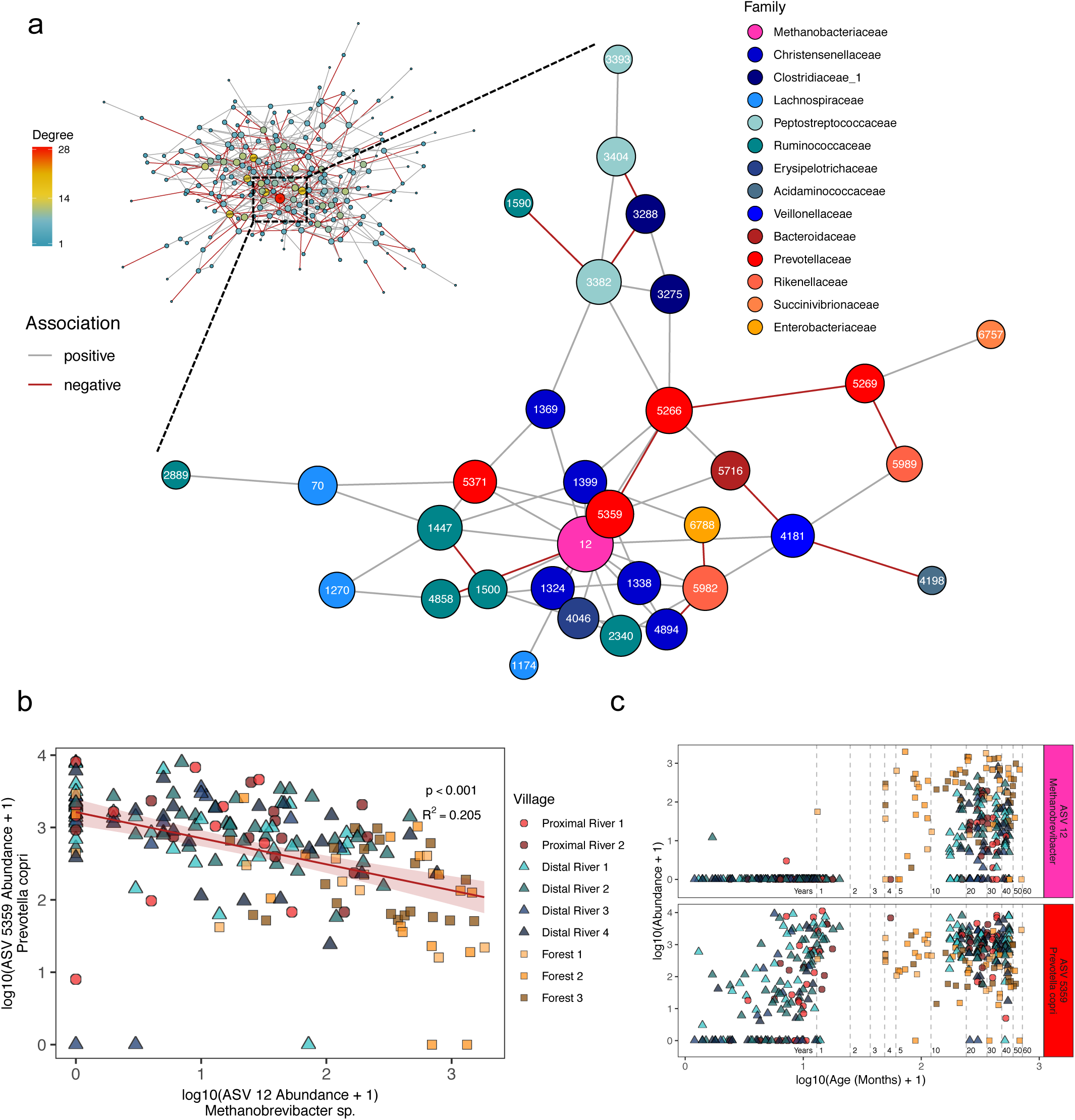
Network analysis reveals an inverse relationship between highly-connected taxa. (a) A network diagram of ASVs observed at least 10 times in at least 20% of Tsimane adult stool samples. The size of the node was scaled to represent its degree of connectedness. The insert shows the complete network colored by network degree, while the main figure shows the highly-connected subgraph colored by each ASV’s taxonomic family. Positive associations between ASV nodes are colored gray, while negative associations are colored red. (b) A scatter plot of the log transformed relative abundances of the two most highly connected ASVs in the network, *Methanobrevibacter* sp. (ASV 12) and *Prevotella copri* (ASV 5359). The red line indicates the linear mixed-effects regression while treating subject as a random effect, and shading indicates the 95% confidence interval. The conditional R^2^ describes the proportion of variation explained by both the fixed and random factors. Samples are colored by village, and their shapes represent the village type (proximal river – circles, distal river – triangles, forest – squares). (c) Scatter plot of the relative abundances of *Methanobrevibacter* sp. (ASV 12) and *Prevotella copri* (ASV 5359) and the subject’s age in months at the time of collection. The x-axis was log transformed for clarity when plotting both infant and adult samples. Ages 1, 2, 3, 4, 5, 10, 20, 30, 40, 50, and 60 years are denoted by horizontal dashed lines.

## DISCUSSION

Here we demonstrate that the Tsimane, an indigenous forager-horticulturalist population, display a distinct microbiota maturation trajectory among their infants. These colonization dynamics may be the result of intensive ICABs. Strain-resolved investigations of mother-infant transmission have found that most vertically transmitted microbes likely originate from the same body site in the mother, and that colonization of strains originating from other maternal body sites is relatively rare^44^. However, we observed evidence of maternal oral-to-infant gut transmission among Tsimane participants, which could be a reflection of traditional intensive ICABs like frequent premastication, and consumption of chicha inoculated with maternal oral microbes. A study of Italian 4-month olds found that 9.5% of bacterial strains in their guts were also found on their mother’s tongue^44^, whereas we observed an average of 23.7% of 3-5 month old Tsimane infant gut ASVs on their mother’s tongue. While metagenomic sequencing allows investigators to track the transmission of specific strains, our 16S rRNA profiling approach, paired with the inference of exact sequence variants, enabled us to perform broad surveys of assembly patterns that can serve as an “upper bound” for estimating transmission rates. This approach also allowed us to apply statistical models to quantify underlying ecological processes that drive host-microbe biology. One important caveat is our assumption that microbes that were shared between mothers and infants imply mother-to-infant transmission. Importantly, we cannot directly observe the direction of transmission, nor can we exclude the possibility that both mothers and children were colonized by microbes from other, unobserved sources.

The patterns of microbial prevalence and abundance fit the NCM less well in both adults and infants living in proximal river villages than did those from villages with less market integration. This suggests that the structure of the Tsimane microbiota may be changing in the face of rapid socioeconomic shifts in ways that decrease the contribution of neutral processes during childhood microbiota assembly, potentially altering interactions between highly abundant microbes like *Methanobrevibacter* and *Prevotella copri.* Perhaps it is not surprising that differences in subsistence strategies have been associated with differences in the gut microbiota of indigenous populations^45^, since even relatively modest differences in lifestyle can also lead to differences in the gut microbiota^46^. A recent study identified microbes that were differentially abundant in indigenous Bassa infants living in rural Nigeria versus infants living in nearby urban centers^47^, hinting that the observed microbiota differences in adults from traditional societies may be influenced by early life factors^40^. Here, we suggest that the differences observed between infants living in traditional societies and those living in or near industrialized cities might be the result of reduced neutral dispersal of microbes that accompanies the hygienic, medical, or societal changes associated with increased market integration. Alternatively, these changes may be the result of maternal microbiotas that have a decreased ability to stably colonize infants. Our findings lend support to the hypothesis that familial microbial transmission in western societies deviates from traditional modes of transmission^48^.

While there are substantial differences in bacterial taxa between adult populations from different regions and cultures, the prominent role of neutral assembly in infant guts is strikingly consistent. This is especially important in the transmission of commensal microbes in indigenous populations^49^, since these relatively isolated societies experience low levels of microbial dispersal from outside their local communities, and exhibit a high degree of homogeneity in their microbiota composition^33^. Collectively, these observations of the microbiota in traditional populations have broad implications for understanding how ecological interactions operate during microbiota assembly, as well as how those patterns are altered due to broad societal changes that have arisen in concert with increased participation in market economies.

## METHODS

### Ethics Approval

All study protocols were approved by the University of California Santa Barbara Institutional Review Board on Human Subjects. Permission to conduct research was granted to the Tsimane Life History Project (THLHP) and their research affiliates. The THLHP maintains formal agreements with the local municipal government of San Borja and the Tsimane governing body. Consent was obtained from village leaders and individual participants upon initiating research activities in each village.

### Sample collection and processing

The two sets of samples (2012-2013 and 2009) were collected as part of a long-term study of Tsimane health and life history, which also included detailed surveys of the subject’s health, diet, and lifestyle. The first set of samples presented in this manuscript was collected from September 2012 through March 2013, and focused on 52 mother-child dyads and used a mixed longitudinal design. These individuals resided in six villages along the Maniqui River. These river villages were grouped into ‘proximal’ or ‘distal’ villages, based on their proximity to the same, nearest market town. Samples were collected together with data on breastfeeding status, 24-hour dietary recall, anthropometrics, and symptoms of infectious disease, as part of a study of changes in infant feeding and related maternal and infant health outcomes^9^. The second analysis presented in this manuscript included samples that were collected in July 2009; a cross-sectional design specified collection of single stool samples from each of 73 Tsimane individuals ranging in age from 1 to 58 years old. These individuals resided in one of three forest villages between 31 and 68 kilometers from the nearest regional market.

Adult and infant fecal samples were collected in plastic containers by the adult subjects, which were then given to investigators typically within 1-2 hours. A portion of each sample was transferred to a cryotube and immediately frozen in liquid nitrogen. Oral samples were collected with a buccal cell collection swab (Catch-All^TM^ Sample CollectionSwab; Epicentre^R^), transferred to cryotubes and stored in liquid nitrogen. All samples were shipped on dry ice to a laboratory in the United States, where they remained at −80°C until they were processed. DNA was extracted from the samples using the MOBIO PowerSoil-htp Kit (MOBIO, Carlsbad, CA) following the manufacturer’s instructions, including a 2 × 10 minute bead-beating step using the Retsch 96 Well Plate Shaker at speed 20. The V4 hypervariable region of the 16S rRNA gene was then PCR-amplified in triplicate using bacterial specific primers (515F and 806R) that include error-correcting barcodes and Illumina adapters^50^. Amplicons were then pooled in equimolar ratios before being sequenced on two runs of an Illumina MiSeq 2×250 PE, generating a total of 35.5 million raw reads.

### 16S rRNA gene sequencing and data processing

Raw reads were denoised into Amplicon Sequence Variants (ASVs) using the DADA2 pipeline (`dada2`, Version 1.9). Following generation of an ASV table, sequences were chimera-checked, and those remaining that were not 230-235 bps in length were removed. In addition to these data, publicly available 16S rRNA datasets from Bangladesh^26^, Finland^27^, and England^35^ were downloaded, run through the DADA2 pipeline, and combined at the level of ASVs with the Tsimane dataset.

Taxonomic assignments were made using the RDP classifier and the SILVA nr database v132. ASVs that were not identified as Domain *Bacteria* or Domain *Archaea* were excluded from further analyses. A phylogenetic tree of the total set of ASVs was inferred using the fragment insertion function (SEPP and `pplacer`) in QIIME2 and the full SILVA v132 tree. An aggregated mapping file from all 4 of the datasets, the ASV table, the taxonomy table, and the phylogenetic tree were imported into R and combined into a single phyloseq object (`phyloseq`, version 1.24.2).

### Data analysis

Both alpha-diversity (Shannon Diversity Index) and beta-diversity (Jaccard Index, Bray-Curtis Distance) analyses were performed in R. Statistical analyses including Wilcoxon tests and PERMANOVA tests were also performed in R. PCoA and canonical correspondence analysis (CCA) ordinations, boxplots, and scatter plots, were generated using the `ggplot2` package in R (version 3.2.0). Heatmaps were generated using the `pheatmap` R package (version 1.0.12). Amplicon sequence variant (ASV) abundances were centered and log ratio-transformed before performing a PCoA on the Jaccard pairwise distances. The partial CCA was performed on centered and log transformed ASV abundances constrained against a matrix of diet survey data, including the 7 MDD food groups, as well as information on meal frequency, frequency of consuming premasticated foods and liquids, frequency of consuming chicha, as well as chicha type. The effects of infant age and village were controlled using a conditioning matrix. Significance was assessed using an ANOVA-like permutation test with 999 permutations. The principal curve was fit to the PCoA in **Fig. 4a** using the `princurve` R package (version 2.1.3).

For the treeDA analysis in **Fig. 5**, ASVs were removed if they were not observed at least 10 times in at least 20% of Tsimane adult stool samples. ASVs were inverse hyperbolic sine-transformed before running the tree-based LDA algorithm in the treeDA R package (version 0.0.3). The LDA score of each adult stool sample was calculated from a phylogeny-based form of linear discriminant analysis optimized with 10-fold cross validation.

Network analysis was performed using the SPIEC-EASI function in the `SpiecEasi` R package (version 1.0.6) to infer an underlying graphical network model using both sparse neighborhood and inverse covariance selection^51^. ASVs were removed that were not observed at least 10 times in at least 20% of Tsimane adult stool samples. The network was constructed with the SPIEC-EASI network inference algorithm using the Meinshausen-Buhlmann Neighborhood Selection method. The size of the nodes were scaled to represent their degree of connectedness.

### Modeling neutral community assembly

As applied by Sloan *et al.* (2006), neutral assembly theory assumes that all microbes in a regional species pool have an equal ability to disperse to a local area, and once established, all have equal fitness, growth, and death rates^34^. These assumptions can be tested using a nonlinear least squares model available in the `minpack.lm` R package (version 1.2-1) to predict the prevalence of a microbe in a local community based on its average relative abundance in the regional species pool. Microbial distributions that are consistent with the model’s predictions are well explained by neutral dispersal, while taxa that deviate from the model likely experience either positive or negative selection. Our approach was a modified version of one used in Burns *et al.*^7^; modeling and plotting functions are available in the `tyRa` R package (version 0.1.0) available at https://danielsprockett.github.io/tyRa/.

### Data and Workflow Availability

Raw sequencing data will be made available at NCBI Sequence Read Archive (SRA Project ID pending). A detailed description of these analyses, along with all analysis code and raw data, are available at the Stanford Digital Repository (https://purl.stanford.edu/tv993xn7633) and as Supplementary Material.

## ACKNOWLEDGEMENTS

We would like to thank the Tsimane host villages and families that participated in this study, as well as the THLHP staff and researchers who provided invaluable assistance during field data collection. We would also like to thank David Sela at the University of Massachusetts Amherst for help with initial study design, and Alvaro Hernandez at the University of Illinois Roy J. Carver Biotechnology Center for outstanding DNA sequencing services. This research was supported by the National Science Foundation NSF BCS 0422690 (M.G.), DDIG 1232370 (M.M.), GRF DGE-114747 (D.S.), National Institute of Health grants NIH/NIGMS T32GM007276 (D.S.), NIH/NIA R01AG024119 (M.G.), NIH/NICHD K99HD074743 (E.K.C.), the Thomas C. and Joan M. Merigan Endowment at Stanford University (D.A.R.), and the Chan Zuckerburg Biohub Microbiome Initiative (D.A.R.).

---

**Supplementary Figure 1.**
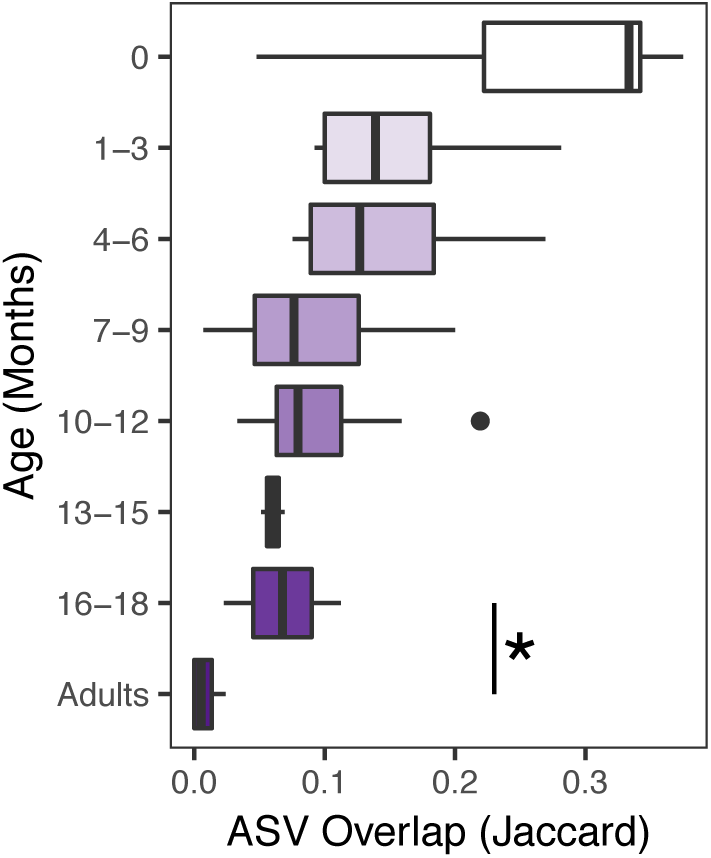
Gut and oral microbiotas diverge as infants age. Average number of ASVs shared between body sites within a subject based on samples collected within 14 days of each other. Boxplot color darkens with increasing age. Adults and 16-18 month olds were the only sequential age groups whose microbiotas were significantly different from one another (Wilcoxon rank-sum test, p = 0.03).

**Supplementary Figure 2.**
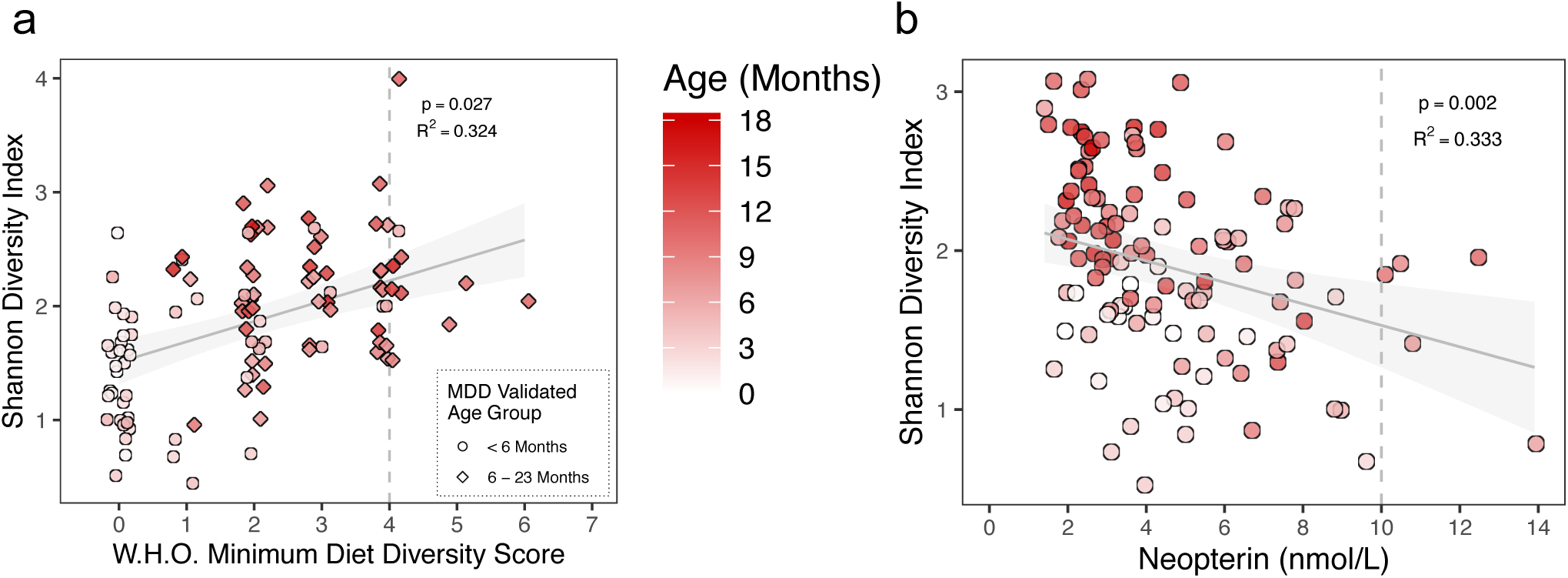
Diet and inflammation impact the infant gut microbiota. (a) The World Health Organization’s minimum diet diversity (MDD) score regressed against the Shannon Diversity Index from the infant stool samples. The MDD was validated to assess the diversity of complementary foods of 6-23 month old children in a diverse range of cultural contexts. It should only be interpreted as an estimate of micronutrient sufficiency in children aged 6-23 months, which are denoted by a diamond shape, and the color darkens as the subject’s age increases. MDD of 4 is considered to be sufficiently diverse, as indicated by the vertical dashed line. Lines indicate the linear mixed-effects regression of diversity on MDD, while treating subject as a random effect. The shading indicates the 95% confidence interval. The conditional R^2^ describes the proportion of variation explained by both fixed and random factors. (B) Levels of fecal neopterin, an inflammatory marker normally produced by activated macrophages, regressed against the Shannon Diversity Index in infant stool samples. Lines indicate the linear mixed-effects regression of diversity on neopterin, while treating subject as a random effect. The shading indicates the 95% confidence interval. The conditional R^2^ describes the proportion of variation explained by both fixed and random factors. A neopterin level above 10, as indicated by the vertical dashed line, indicates clinically relevant inflammation.

**Supplementary Figure 3.**
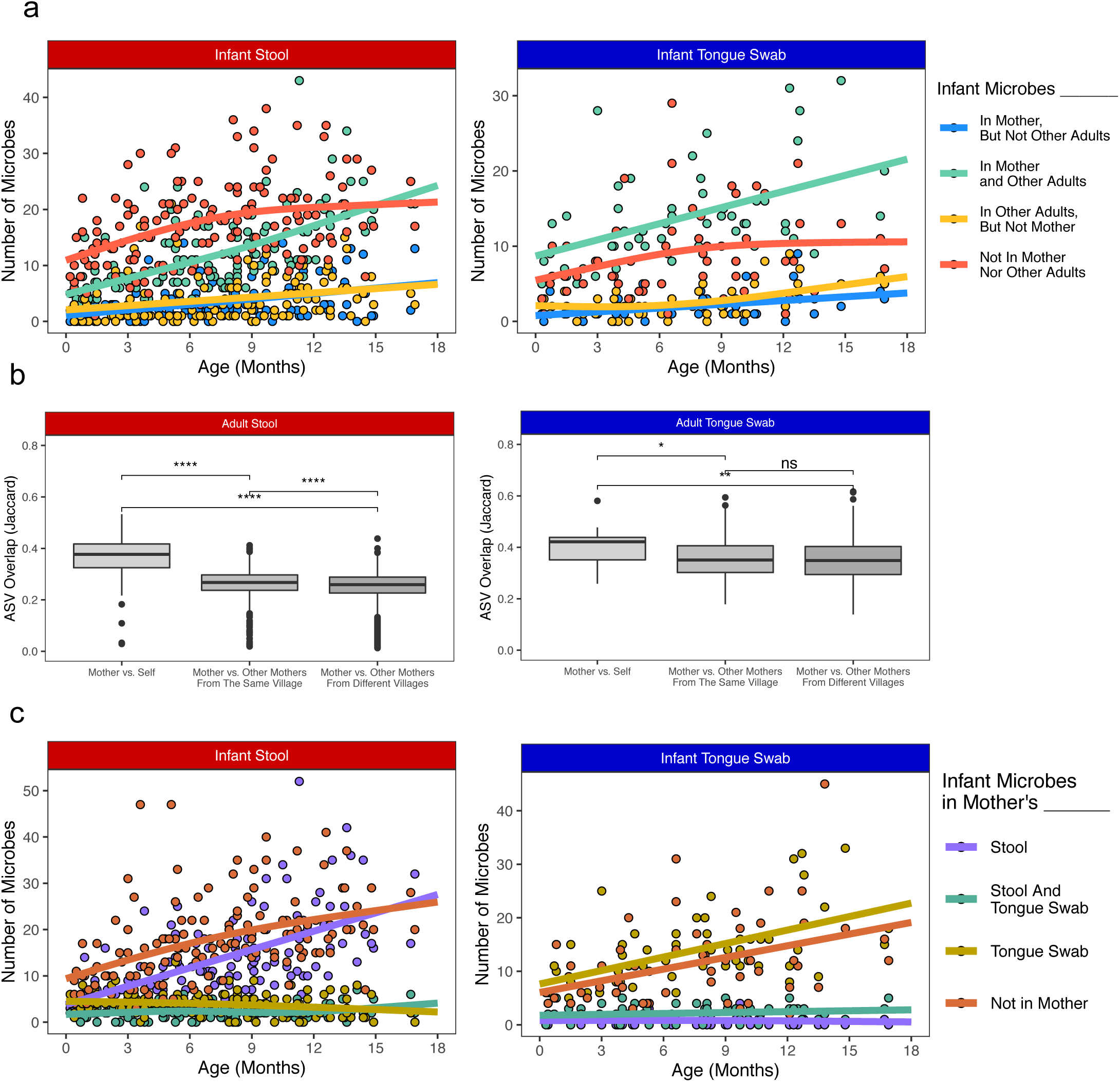
Sources of infant-colonizing microbes. (a) Scatter plot of the average number of ASVs found in each infant stool sample that were also found in that infant’s mother but not in other adults from their village (blue), in both their mother and other mothers from their village (green), in other mothers from the infant’s village but not their mother (yellow), or in neither their mother or other adults (orange). Non-parent adults from each village were down-sampled to control for differences in the number of samples collected from each village. Left panel – infant stool samples, right panel – infant tongue swabs. (b) Average number of ASVs shared between multiple samples from the same mother, between mothers living in the same village, or mothers living in different villages. Wilcoxon Rank Sum Test. ****: p ≤ 0.0001, **: p ≤ 0.01, *: p ≤ 0.05, ns: p > 0.05. Left panel – adult stool samples, right panel – adult tongue swabs. (c) Scatter plot of the average number of ASVs in each infant stool sample that was also found in their mother’s stool but not in her tongue swab (purple), in both their mother’s stool and tongue swab (green), in their mother’s tongue swab but not their mother’s stool (yellow), or not found in any of their mother’s samples (orange).

**Supplementary Figure 4.**
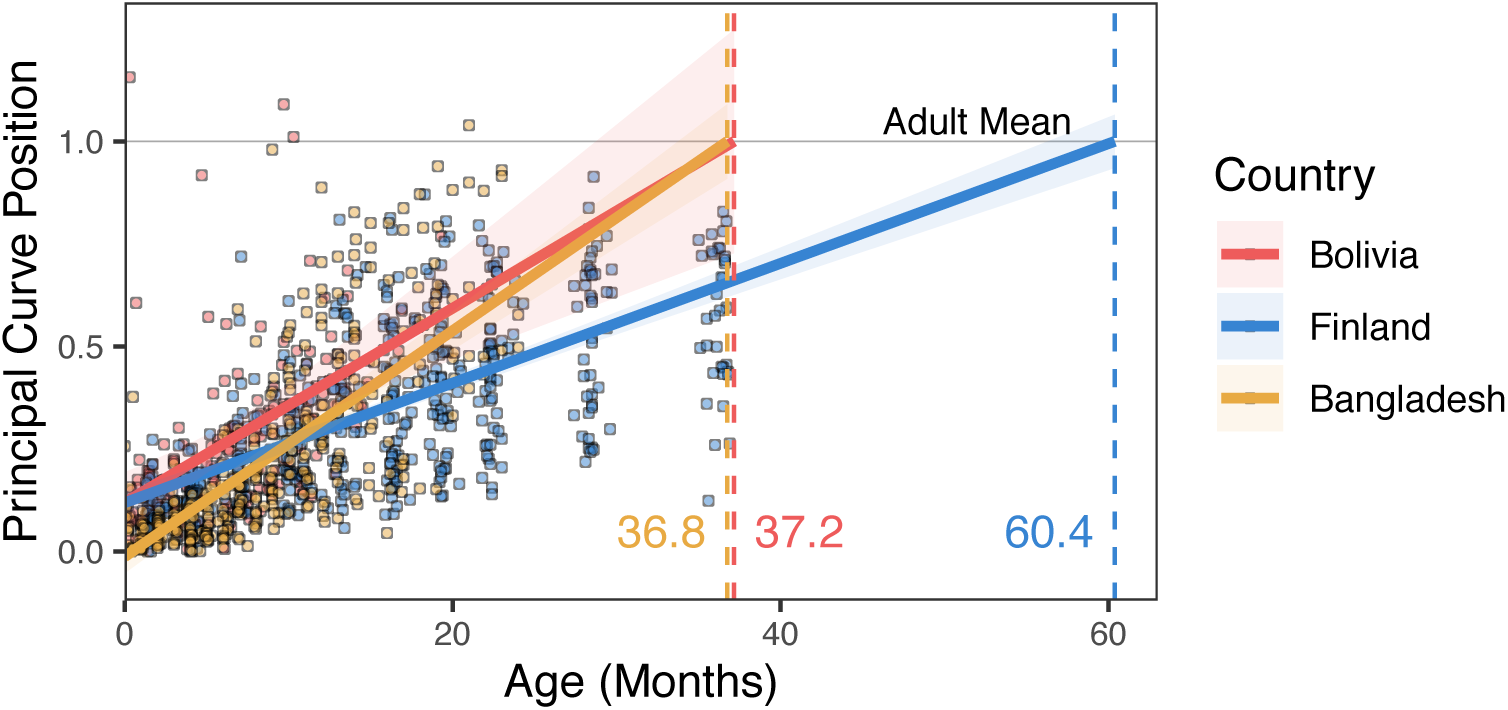
Infants from different countries reach a population-specific adult-like state at different rates. (a) In Fig. 4a, a PCoA was performed on 16S rRNA datasets from three populations (Bolivian (Tsimane), Bangladeshi, and European (Finnish/ British)), and a principal curve was fit to each set of points using the princurve function in the princurve R package (version 2.1.3). The distance along each curve was scaled so that the mean location of the adult samples was equal to one (as indicated by the horizontal gray line), making an infant’s location along the curve analogous to their position along the average maturational trajectory towards the average adult in their society. Here, we show the linear regression of the age of the infant from whom each sample was collected against its position along its principal curve. Lines indicate the linear mixed-effects regression of principal curve position on the infant’s age in months, while treating subject as a random effect. The shading indicates the 95% confidence interval. The vertical dashed lines indicate the age at which the infant line from each country crosses the adult average position for that country.

**Supplementary Figure 5.**
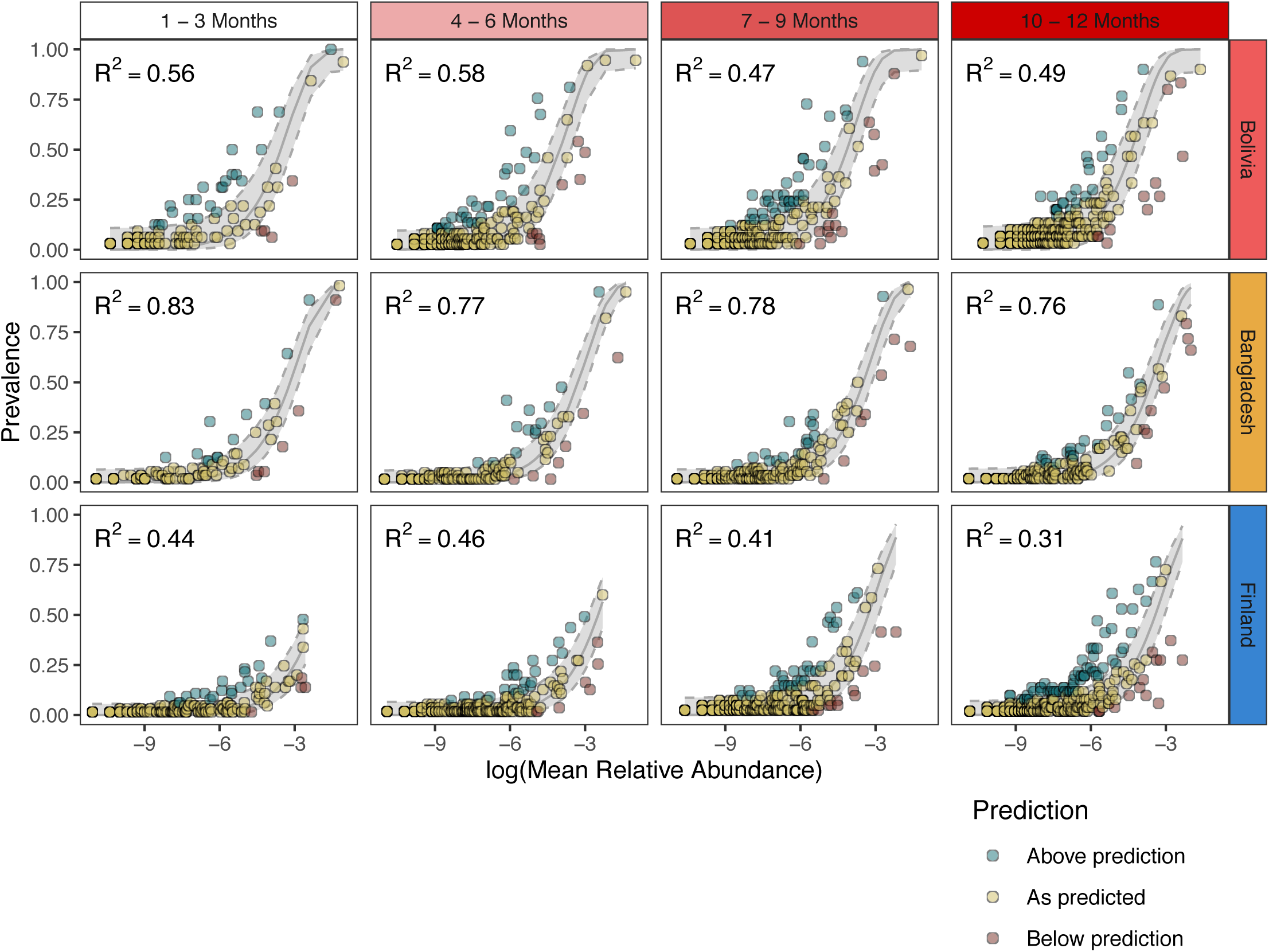
Neutral processes explain microbial community assembly across age groups and geography. Scatterplot of the prevalence of each ASV in the infant gut versus its mean relative abundance in the regional species pool for each age group and country. The gray line is their predicted distribution (shaded area is 95% confidence interval) based on the neutral community model. Points are colored by the ASV’s fit to the model: above prediction– green, below prediction– red, at prediction – yellow.

**Supplementary Figure 6.**
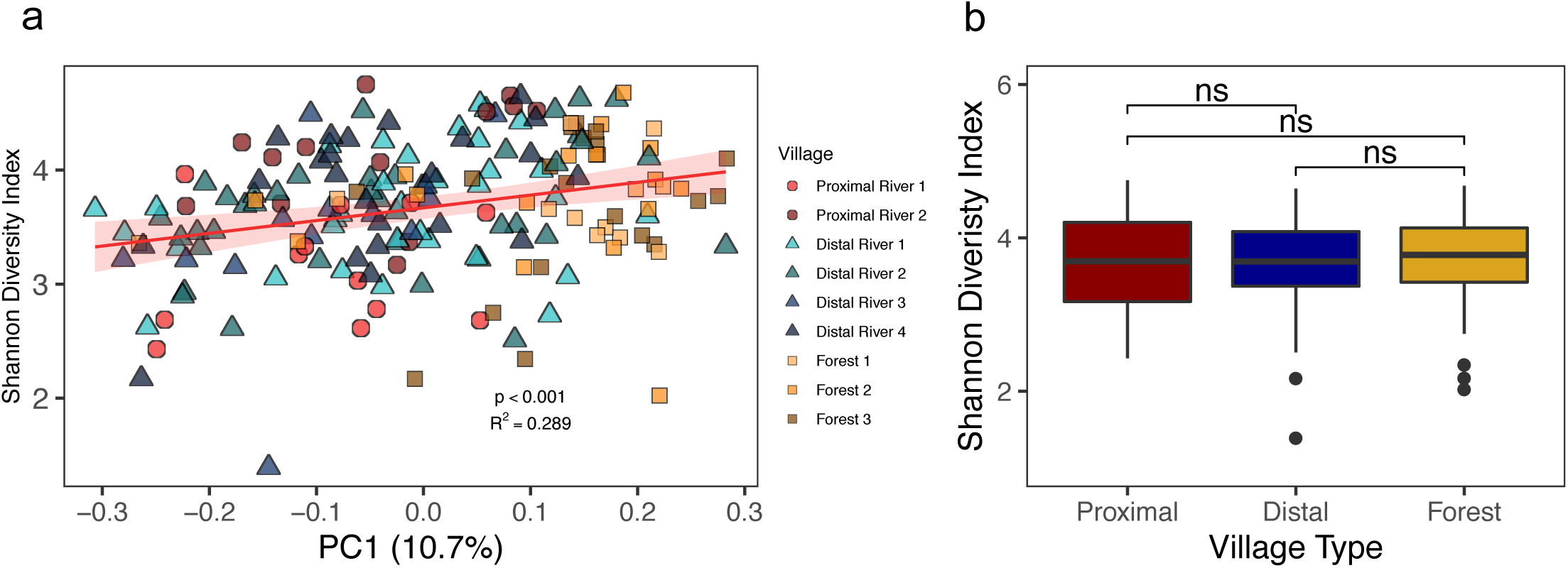
Gut microbiota diversity among Tsimane villages. (a) Shannon Diversity Index plotted against the PC1 value from Fig. 5b (R^2^ = 0.289, p < 0.001). The red line indicates the linear mixed-effects regression while treating subject as a random effect, and shading indicates the 95% confidence interval. The conditional R^2^ describes the proportion of the variation explained by both fixed and random factors. (b) Boxplot comparing the average diversity of adult stool microbiota of samples collected from proximal river, distal river, and forest Tsimane villages (p > 0.05 for all comparisons).

**Supplementary Figure 7.**
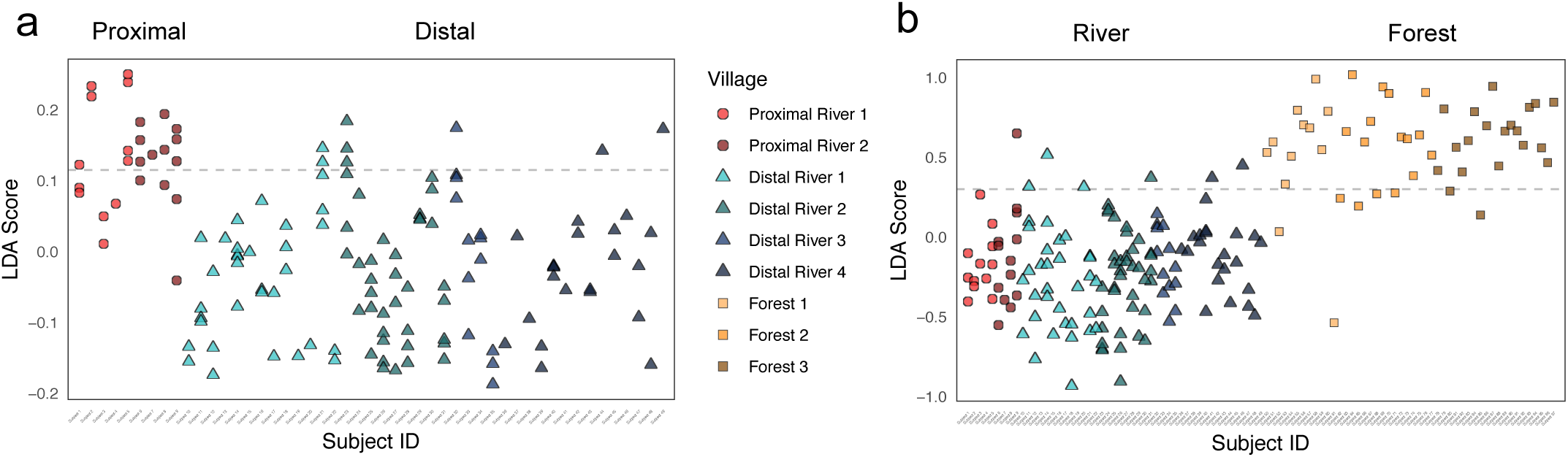
Bacterial taxa discriminate between village types. (a) The LDA score of each adult stool sample was calculated from a phylogeny-based form of linear discriminant analysis (treeDA, version 0.0.3). Higher scores indicate samples more likely to have been collected from a proximal river village, while lower scores indicate samples more likely to have been collected from a distal river village. The dashed gray line represents the optimal LDA score cut-off between groups, where it accurately classifies 87.3% of samples. Samples are colored by village, and their shapes represent the village type (proximal river – circles, distal river – triangles, forest – squares). (b) The LDA score of each adult stool sample was calculated from a phylogeny-based form of linear discriminant analysis (treeDA, version 0.0.3). Higher scores indicate samples more likely from a forest village, while lower scores indicate samples more likely from a river village. The dashed gray line represents the optimal LDA score cut-off between groups, where it accurately classifies 91.8% of samples. The colors and shapes are the same as in panel a.

**Supplementary Table 1.** Demographics of Tsimane villages and study subjects. Villages are listed in order of increasing distance from the closest market town. Age classes are defined as Infant – under 2 years, Child – 2 to 13 years old, Adult – 14 years old and older. The 73 fecal samples collected from forest communities were collected in July 2009, while the remaining fecal and oral samples from proximal and distal river communities were collected between September 2012 and March 2013. *Population sizes for forest villages were collected during a 2009 census, while the remaining were collected in a 2012 census.

| Village | Age Class | Number of Subjects | Stool Samples | Tongue Swabs | All Samples | Distance from City (Km) | Population* |
| --- | --- | --- | --- | --- | --- | --- | --- |
| Proximal River 1 | *Adult* | 6 | 12 | 8 | 47 | 14.3 | 331 |
|  | *Child* | 0 | 0 | 0 |  |  |  |
|  | *Infant* | 6 | 20 | 7 |  |  |  |
| Proximal River 2 | *Adult* | 4 | 13 | 6 | 42 | 18.5 | 292 |
|  | *Child* | 0 | 0 | 0 |  |  |  |
|  | *Infant* | 4 | 17 | 6 |  |  |  |
| Distal River 1 | *Adult* | 13 | 34 | 13 | 92 | 29.8 | 151 |
|  | *Child* | 0 | 0 | 0 |  |  |  |
|  | *Infant* | 11 | 33 | 12 |  |  |  |
| Distal River 2 | *Adult* | 9 | 39 | 18 | 128 | 33.5 | 253 |
|  | *Child* | 0 | 0 | 0 |  |  |  |
|  | *Infant* | 10 | 53 | 18 |  |  |  |
| Distal River 3 | *Adult* | 5 | 13 | 5 | 36 | 37.5 | 101 |
|  | *Child* | 0 | 0 | 0 |  |  |  |
|  | *Infant* | 5 | 14 | 4 |  |  |  |
| Distal River 4 | *Adult* | 14 | 23 | 12 | 65 | 42.5 | 206 |
|  | *Child* | 0 | 0 | 0 |  |  |  |
|  | *Infant* | 12 | 19 | 11 |  |  |  |
| Forest 1 | *Adult* | 12 | 12 | 0 | 21 | 31 | 61 |
|  | *Child* | 8 | 8 | 0 |  |  |  |
|  | *Infant* | 1 | 1 | 0 |  |  |  |
| Forest 2 | *Adult* | 16 | 16 | 0 | 27 | 61 | 114 |
|  | *Child* | 11 | 11 | 0 |  |  |  |
|  | *Infant* | 0 | 0 | 0 |  |  |  |
| Forest 3 | *Adult* | 20 | 20 | 0 | 25 | 68 | 101 |
|  | *Child* | 5 | 5 | 0 |  |  |  |
|  | *Infant* | 0 | 0 | 0 |  |  |  |

